# Cardiolipin-Dependent Properties of Model Mitochondrial Membranes from Molecular Dynamics Simulations

**DOI:** 10.1101/557744

**Authors:** Blake A. Wilson, Arvind Ramanathan, Carlos F. Lopez

**Affiliations:** Department of Biochemistry, Vanderbilt University, 2201 West End Ave, Nashville, TN 37235; Computational Science and Engineering Division, Oak Ridge National Laboratory, P.O. Box 2008 Oak Ridge, TN 37831; Health Data Sciences Institute, Oak Ridge National Laboratory, P.O. Box 2008 Oak Ridge, TN 37831; Department of Biomedical Informatics, Vanderbilt University Medical Center, 1211 Medical Center Drive, Nashville, TN 37232; Department of Pharmacology, Vanderbilt University, 2201 West End Ave, Nashville, TN 37235

## Abstract

Cardiolipin is a unique anionic lipid found in mitochondrial membranes where it contributes to various mitochondrial functions, including metabolism, mitochondrial membrane fusion/fission dynamics, and apoptosis. Dysregulation of cardiolipin synthesis and remodeling have also been implicated in several diseases, such as diabetes, heart disease and Barth Syndrome. Although cardiolipin’s structural and dynamic roles have been extensively studied in binary mixtures with other phospholipids, the biophysical properties of cardiolipin in ternary lipid mixtures are still not well resolved. Here, we used molecular dynamics simulations to investigate the cardiolipin-dependent properties of ternary lipid bilayer systems that mimic the major components of mitochondrial membranes. We found that changes to cardiolipin concentration only resulted in minor changes to bilayer structural features, but that the lipid diffusion was significantly affected by those alterations. We also found that cardiolipin position along the bilayer surfaces correlated to negative curvature deflections, consistent with the induction of negative curvature stress in the membrane monolayers. This work contributes to a foundational understanding of the role of CL in altering the properties in ternary lipid mixtures composed of the major mitochondrial phospholipids, providing much needed insights to help understand how cardiolipin concentration modulates the biophysical properties of mitochondrial membranes.

## 1 INTRODUCTION

Cardiolipin (CL) is an anionic lipid found in the membranes of eukaryotic mitochondria (1), as well as the inner membrane of chloroplasts and some bacterial plasma membranes. The signature mitochondrial lipid, CL contributes to various mitochondrial functions including metabolism (2), mitochondrial membrane fusion/fission dynamics (3), and apoptosis (4, 5). Furthermore, dysregulation of CL synthesis and remodeling has been implicated in several diseases, among them diabetes (6, 7), heart disease (7, 8), and Barth Syndrome (9, 10).

Mitochondrial CL is primarily located in the inner mitochondrial membrane (IMM) (11–16) where it accounts for ∼10 mol% of IMM lipid content. CL molecules bind promiscuously to IMM proteins (1, 17), stabilizing various enzymes and protein complexes involved in oxidative phosphorylation (2, 18) which in turn facilitates optimal metabolic energy conversion. There is also evidence that CL plays a role in maintaining IMM superstructure (19–23), affecting the shape and stability of cristae.

CL is also present in the outer mitochondrial membrane (OMM) (11–16), but at lower concentrations (∼ 1 – 5 mol% of OMM lipids) and with higher concentration fluctuations than in the IMM. The OMM is both the ‘locus of action’ and an active participant in the interactions that regulate the mitochondrial apoptosis pathway (24–26). Execution of the mitochondrial apoptosis pathway culminates with permeabilization of the OMM, an event that is tightly regulated by a complex network of interactions between members of the Bcl-2 family of proteins and the OMM (27, 28). CL contributes to this process by acting primarily as an accelerator of tBid-mediated MOMP, facilitating the recruitment of the protein to the membrane and the activating conformational changes that allow tBid to efficiently recruit and activate the pore forming Bcl-2 proteins Bax/Bak (26, 29–35). Additionally, CL was shown to affect the kinetics of both tBid-Bax and Bim-Bax heterodimer interactions (26) and to enhance the permeabilization function of activated oligomerized Bax (in the absence of tBid) (30), suggesting that CL may modulate other interactions in the MOMP regulatory network. CL content has also been correlated with preferential localization of tBid to OMM contact sites with the IMM (29, 36), which are locally enriched in CL (up to ∼25 mass%) (16). In addition, it has been shown that CL concentration increases in the OMM during apoptosis execution (37), thus raising questions about how CL concentration modulates OMM properties.

CL is a structurally unique dimeric phospholipid that is composed of two phosphatidylglycerol moieties connected by a single bridging glycerol group (38). CL therefore has a small polar headgroup with limited flexibility and mobility. The small size of the polar headgroup imparts CL with a propensity to form inverted hexagonal lipid phases (39, 40), while many of CL’s unique membrane properties have been attributed to the restrictions placed on the polar headgroup’s flexibility and mobility (41).

The two major phospholipid species in mitochondrial membranes are phosphatidylcholine (PC) and phosphatidylethanolamine (PE) lipid species. A variety of studies, including monolayer experiments (42–45), vesicle and bilayer experiments (42, 46–48), and simulation (49–51) have probed the biophysical properties of binary mixtures of CL with PC or PE lipids. A limited number of experimental studies have also probed the biophysical properties of more complex lipid mixtures (>ternary) using OMM-like supported lipid bilayers (48) and IMM-like monolayers (45). However, the studies that investigate the properties of CL in mixed PC-PE lipid membranes are limited, so there is still a shortage of basic biophysical data on CL in ternary lipid membrane systems.

The main purpose of this work is to investigate the CL-dependent properties of ternary lipid bilayer systems that mimic the major components of mitochondrial membranes. There is ample evidence that membrane material properties (lipid packing, thickness, fluidity, etc.) play key roles in the function of membrane proteins, affecting their conformation (52–56), activity (57–60), and distribution (61, 62). These properties modulate the landscape of interactions primarily through their effects on the energy and kinetics of protein-membrane and protein-protein interactions at the membrane (63, 64). Evaluating the CL-dependent properties of model mitochondrial membranes can help elucidate CL’s role in the structural differences between the IMM and OMM. And the CL-dependent properties are also particularly relevant for understanding how changes in mitochondrial CL content, such as during the induction of apoptosis (37), may have on the mitochondrial membranes. As such, we have run a series of atomistic molecular dynamics (MD) simulations of model mitochondrial membranes with CL concentrations consistent with the range of CL content found in the OMM, IMM, and their contact sites in the OMM. From these simulations we have estimated a variety of structural, mechanical, and dynamic bilayer properties and show their dependence on the concentration of CL. We also compare our results to those of previous studies of the thermodynamic, structural, and dynamic properties of similar ternary PC, PE, and CL lipid membranes.

## 2. METHODS

### 2.1 Model Systems

We have constructed five atomistic models of ternary lipid bilayer systems mimicking mitochondrial membranes. The membrane bilayers were square patches with sizes of ∼(14-15 nm)^2^ and were composed of PC, PE and varying proportions of CL molecules consistent with those found in the outer (∼1 – 5 mol%, ∼ 15 mol% at contact sites) and inner (∼ 5 – 10 mol%) mitochondrial membranes. The classes of lipids were modeled using the following lipids:

1. PC: 1-palmitoyl-2-oleoyl-sn-glycero-3-phosphocholine (POPC)
2. PE: 1,2-dioleoyl-sn-glycero-3-phosphoethanolamine (DOPE)
3. CL: 1,3-Bis-[1,2-di-(9,12-octadecadienoyl)-sn-glycero-3-phospho]-sn-glycerol (tetralinoleoyl cardiolipin, (18:2)_4_-CL)

We chose polyunsaturated tetralinoleoyl CL ((18:2)_4_-CL) due to its high relative abundance among cardiolipin species in human mitochondria, particularly in heart and skeletal muscle tissues (65–67). Tetralinoleoyl CL is also the primary cardiolipin species affected in Barth Syndrome (9, 66).

Since they have two phosphate groups, CL molecules can carry up to two negative charges. However, the ionization state of cardiolipin at neutral pH (7.0) and physiological pHs (around pH 7.4) is a contentious issue, with some studies (68–70) suggesting that a mixture of cardiolipin ionization states predominated by the singly-deprotonated (−1e charge) species would likely form, whereas other studies (71–73) suggest that CLs would be fully deprotonated (−2e charge) at that pH range. Since even in the former case the fully ionized CL species may be expected to be present in reasonable proportion, we modeled CL lipids as fully ionized. The structures of the lipids used in our computational models are shown in Figure 1.

**Figure 1:**
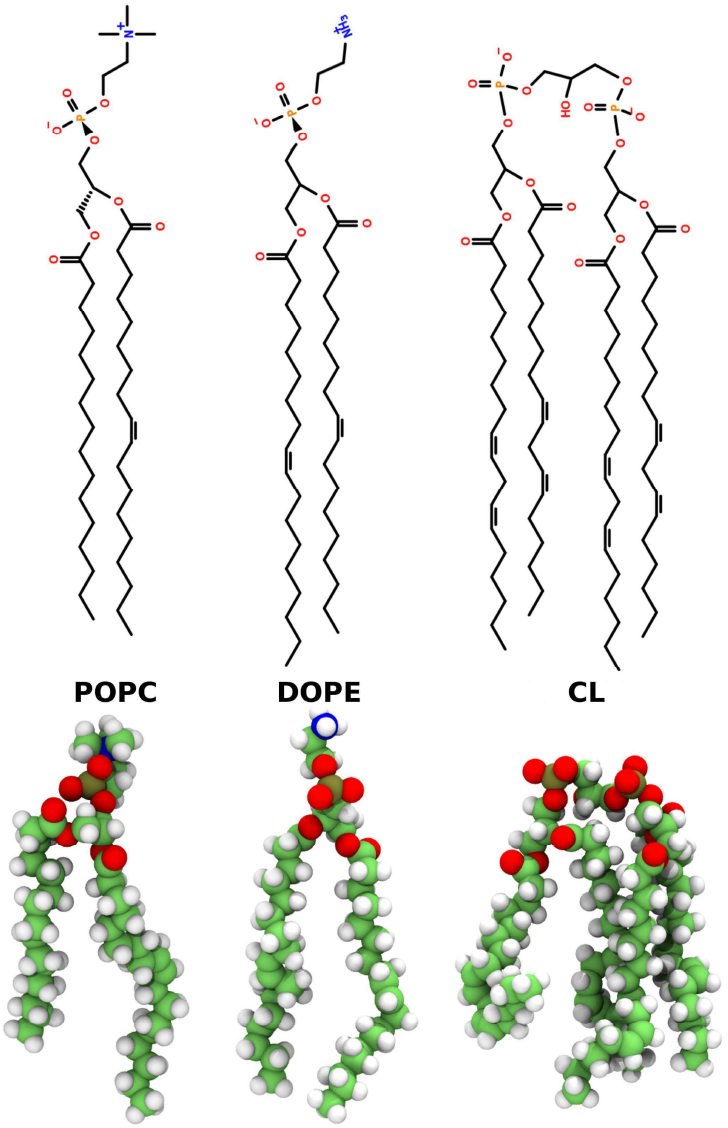
The line (top) and 3-d structures (bottom) of the lipids 1-palmitoyl-2-oleoyl-sn-glycero-3-phosphocholine (POPC), 1,2-dioleoyl-sn-glycero-3-phosphoethanolamine (DOPE), and the −2e charge version of the lipid tetralinoleoyl cardiolipin (CL) used in the model mitochondrial outer membranes. The line structure drawings were generated using Lipid MAPS [http://www.lipidmaps.org/] and Open Babel [http://openbabel.org] tools (74, 75). The 3-d structure images were generated using the Tachyon renderer [http://jedi.ks.uiuc.edu/∼johns/raytracer/] in VMD [http://www.ks.uiuc.edu/Research/vmd/] (76, 77).

All-atom lipid bilayers were constructed using the Membrane Builder Input Generator of the CHARMM-GUI web toolkit [http://www.charmm-gui.org/] (78–83). Each MOM-like bilayer system consisted of 600 lipids (300 per leaflet) solvated with 22.5 of explicit water molecules on either side of the bilayer (37-43 water molecules per lipid). We generated membranes with CL content 0, 2, 7, 10, and 15 mol% (0, 3.8, 12.6, 17.6, and 25.3 mass% respectively); the total number of lipids (600) was held constant. The numbers of POPC and DOPE lipids were adjusted such that an approximate 2:1 ratio (POPC:DOPE) was maintained. Sodium chloride ions were added to neutralize excess lipid charges and set the salt concentration to approximately 0.2 M in the 0, 7, 10, 15 mol% systems. The 2 mol% system was only charge neutralized with Na^+^ ions; although this gives the 2 mol% system slightly different ionic conditions than the other model systems, based on previous studies (84, 85) we expect this difference in ion conditions to have a negligible effect on the properties reported in this study. Interactions were modeled using the CHARMM36 forcefield (86) with the TIP3P water model (87). Representations of the model systems are depicted in Figure 2

**Figure 2:**
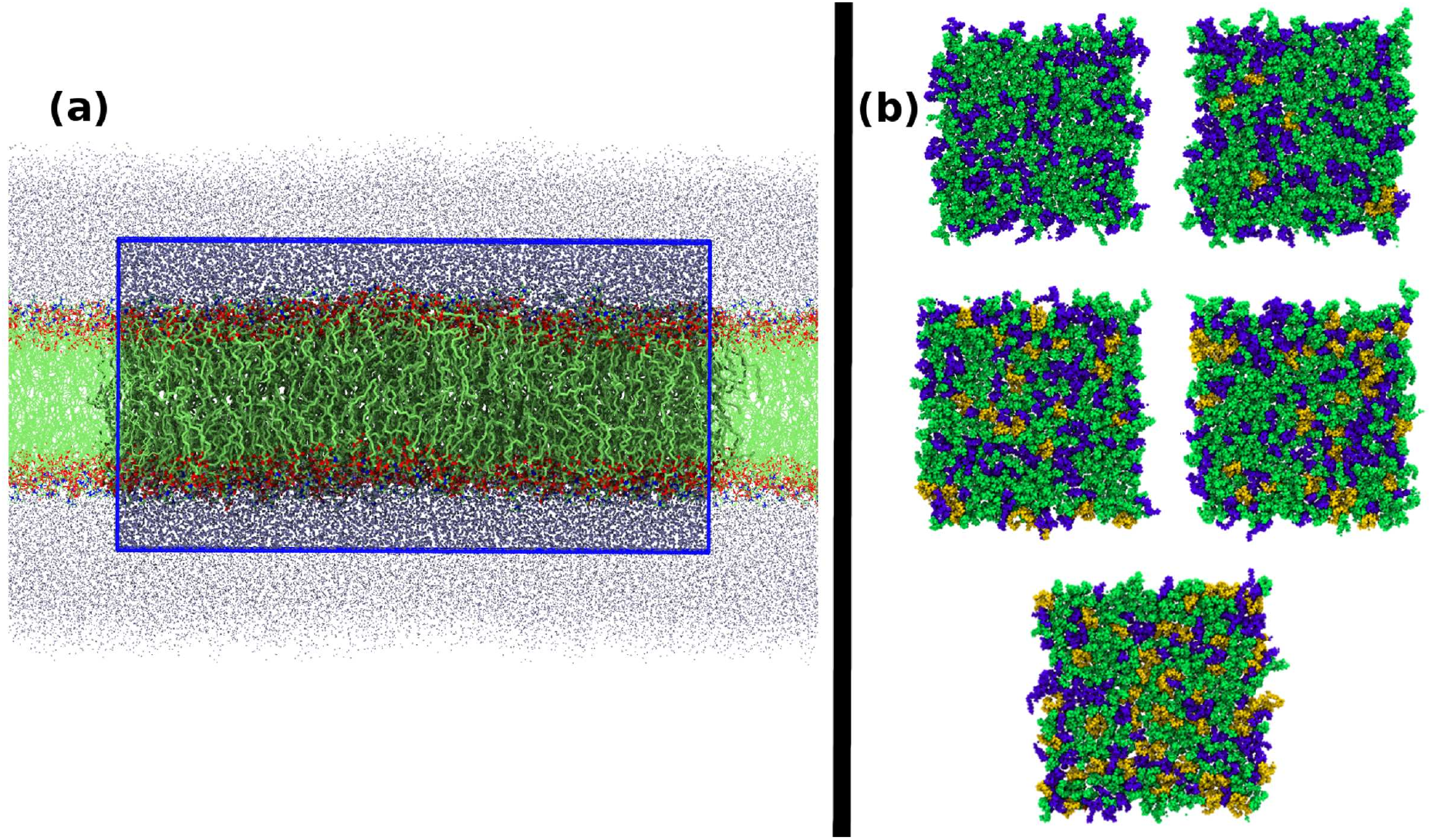
Snapshots of lipid bilayers used to model mitochondrial outer membranes. **(a)** Representative side-view of one of the model systems. For clarity the hydrogen atoms were removed. Water oxygen atoms are shown in ice-blue. The simulation cell is outlined by the blue rectangular box and the material surrounding the simulation cell shows the periodic images. **(b)** Top-view of each model bilayer after initial system minimization and relaxation. The CL concentrations are in increasing order going from left to right and top to bottom. POPC lipids are shown in green, DOPE lipids in blue, and CL lipids in gold. The images of simulation snapshots were generated using the Tachyon renderer [http://jedi.ks.uiuc.edu/∼johns/raytracer/] in VMD [http://www.ks.uiuc.edu/Research/vmd/] (76, 77).

### 2.2 Molecular Dynamics simulations

Simulations were run employing the GROMACS software [version 5.1.4, http://www.gromacs.org/] (88, 89). The pre-production minimization and relaxation of each system was carried out using the GROMACS protocol generated by the CHARMM-GUI Input Generator, which consists of

1. Energy-minimization for 5,000 steps
2. Constant volume and 303.15 K (NVT) using the Berendsen thermostat for 50,000 steps with a 1 fs timestep
3. 1.0 bar and 303.15 K (NPT) using the Berendsen thermostat and barostat for 25,000 steps with a 1 fs timestep
4. 1.0 bar and 303.15 K (NPT) using the Berendsen thermostat and barostat for 150,000 steps with a 2 fs timestep

The production runs were simulated at 1.0 bar and 303.15 K (NPT) using the Nose-Hoover thermostat and the Parrinello-Rahman barostat with a 2 fs timestep; the 2, 7, and 10 percent CL systems were simulated for a total of 1 µs while the 0 and 15 percent CL systems were simulated for a total of 0.9 µs.

All simulations were carried out with covalent bonds to hydrogen atoms constrained via the LINCS algorithm (90). A 12 Å spherical cutoff was used for short-range non-bonded interactions with a force switching function from 10 Å for the van der Waals term and shifted electrostatics. Long-range electrostatic interactions were computed using the particle-mesh Ewald (PME) method (91) with a grid spacing on the order of 1 Å or less. The equations of motion were integrated using the default GROMACS’ ‘md’ integrator. Periodic boundary conditions were applied to all simulations, and all NPT ensemble simulations were run under zero tension and semi-isotropric pressure coupling.

### 2.3 Analysis

Analysis was performed using tools from PyBILT, which is a Python based lipid bilayer analysis toolkit (https://github.com/LoLab-VU/PyBILT) currently being development in our lab. Descriptions of individual methods for analysis are provided in subsequent sections. In lipid membrane systems the dynamics of some properties, such as ion binding/unbinding to the membrane-water interface (84) and lipid diffusion (92), can have characteristic times on the order of 100 ns or greater. Therefore, the first 400 ns of each production run was discarded as a long time-scale equilibration time and not included in the analysis. Unless otherwise noted, thermodynamic averages and their standard errors were estimated by using the block averages taken over non-overlapping 100 ns segments of the remaining trajectory. Error bars for our data were computed as 1.96 times the standard error estimate (from block averaging) and thus represent estimates of the 95% confidence intervals.

#### 2.3.1 Hexagonal packing order parameter

A hexagonal packing order parameter gauges the extent to which lipids lipids and their nearest-neighbors are packed in a hexagonal geometry. We adapted the approach by Katira et al. (93) for hexagonal packing calculation of lipid acyl chains. Via this approach, we computed the ensemble average of Halperin and Nelson’s local rotational-invariant,

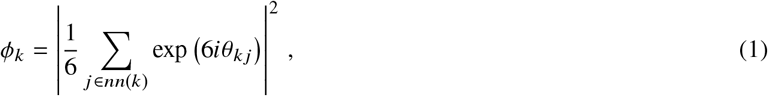

with the terminal carbon atoms of the acyl chains as reference points projected into the xy-plane. In Equation 1, *θ*_*k j*_ is the angle between an arbitrary axis (we used the x-axis) and the 2-d vector in the bilayer lateral plane (xy-plane) connecting the terminal acyl chain carbon *l* to terminal acyl chain carbon *j*. The summation *j ∈ nn k* is taken over the six nearest neighbors (in the bilayer lateral plane) of terminal acyl chain carbon *k*. The ensemble average of *Φ*_*k*_ is unity for a perfect hexagonal packing and zero in the absence of any hexagonal packing.

#### 2.3.2 Average area per lipid molecule

The average area per lipid molecule was computed using the area projected onto the xy-plane, which was taken as the xy simulation box area, divided by the number of lipid molecules per bilayer leaflet:

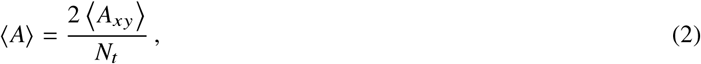

where *A*_*xy*_ is the instantaneous xy-area of the simulation box (used to estimate the lateral surface area of the bilayer) at each simulation snapshot and *N*_*t*_*/*2 is the number of lipids per bilayer leaflet; *N*_*t*_ = 600 was the total number of lipids in the bilayer and was constant for all simulated systems.

#### 2.2.3 Average area per lipid phosphate headgroup

The average area per lipid phosphate headgroup (abbreviated hereafter as average area per phosphate) was estimated by computing the average area per lipid phosphate headgroup projected onto the xy-plane, calculated as the xy simulation box area divided by the number of lipid phosphate headgroups per leaflet:

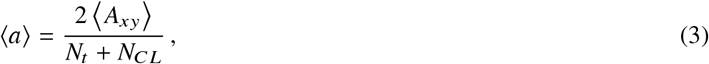

where *A*_*xy*_ is the instantaneous xy-area of the simulation box (used to estimate the lateral surface area of the bilayer) at each simulation snapshot and (*N*_*t*_ + *N*_*CL*_)/2 is the number of lipid phosphate headgroups per bilayer leaflet; *N*_*CL*_ is the total number of CL lipid molecules in the bilayer. Note that this quantity is directly related to the average area per lipid through the conversion factor *N*_*t*_ /(*N*_*t*_ + *N*_CL_); i.e., 〈*a*〉 = 〈*A*〉 *N*_*t*_ /(*N*_*t*_ + *N*_CL_).

#### 2.3.4 Partial area per lipid phosphate headgroup

The partial area per lipid phosphate headgroup was computed using a grid-based tesselation scheme adapted from Allen et al. (94). Briefly, at each snapshot of the simulation trajectory the lipids are mapped to a 2-dimensional grid based on Euclidean distance in the lateral plane of the bilayer (xy-plane). The positions of the phosphate phosphorous atoms projected into the xy-plane as the reference points for distance calculations and subsequent mapping of the lipid headgroups to the 2-d grids. For each grid element the nearest lipid is assigned to that grid. We performed this analysis with 100×100 grids, resulting in grid elements with an area < 3 Å^2^. Gapsys et al. (95) reported that an area per grid element of < 5 Å^2^ is sufficient for convergence of this type of grid based area analysis. A grid map is generated for each leaflet of the bilayer, resulting in two grid maps per analyzed snapshot. The partial area per lipid phosphate headgroup is computed for lipid type *l* from the grids via the following equation,

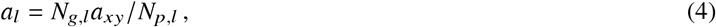

where *N*_*g,l*_ is the total number of grid elements occupied by lipid type *l* (including both leaflets), *a*_*xy*_ = *A*_*xy*_/100^2^ is the area per grid element, and *N*_*p,l*_ is the number of phosphate headgroups that lipid type *l* has in the bilayer. Note that as described, this analysis yields the composite partial areas over both bilayer leaflets and assumes molecules of each lipid type are equally distributed across those leaflets.

#### 2.3.5 Average area per lipid acyl chain

The average area per lipid acyl chain was computed in the same manner as the average area per lipid molecule (Section 2.3.2) or the average area per phosphate (Section 2.3.3), except in this case, the xy simulation box area (*A*_*xy*_) was divided by the number of lipid acyl chains per leaflet. Note that since each lipid has two acyl per chains per phosphate headgroup the average area per lipid acyl chain is equivalent to the average area per phosphate multiplied by a factor of 1/2.

#### 2.3.6 Partial area per lipid acyl chain

The partial areas per lipid acyl chain were computed in the same manner as the partial areas per phosphate (Section 2.3.4), except in this case, the positions of the terminal acyl chain carbon atoms projected into the xy-plane were used as the reference points for distance calculations and subsequent mapping of the lipids to the 2-d grids.

#### 2.3.7 Relative fractions of lipid-lipid interactions

The relative fractions of lipid-lipid interactions were computed using a nearest neighbour analysis as in de Vries et al. (96). For each lipid-lipid interaction of type X-Y, the fraction *f*_*X*-*Y*_ was computed:

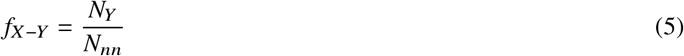

with *N*_*Y*_ the number of lipids of type *Y* amongst the *N*_*nn*_ nearest neighbours of lipid type *X*. We used *N*_*nn*_ = 5 for our calculations, and the positions of the lipids’ centers of mass projected into the xy-plane were used as the reference points for distance calculations and subsequent construction of nearest neighbor lists.

#### 2.3.8 Dimensionless packing shape parameter

The dimensionless packing shape parameter (97), *P* = v/(*l*_*c*_ *a*_*o*_), of each lipid type can be used to quantify lipid shape (98); in that expression, v is the volume occupied by the lipid, *l*_*c*_ is the length of its hydrophobic region, and *a*_*o*_ is the area occupied by its polar headgroup. For our analysis, we assumed each lipid within the bilayer was a truncated cone, cylinder, or inverted truncated cone shape (using shapes as defined in Figure 1 of Dutt et al. (99)). In this case, another parameter for the tail-end area occupied by the lipid acyl chains, *a*_*h*_, can be introduced (98) and the the equation for the packing shape parameter can be rewritten as:

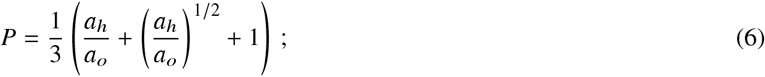

this parameter is equal to 1 for cylinder shaped lipid molecules, <1 for truncated cone shaped lipid molecules, and >1 for inverted truncated cone shaped lipid molecules. For each lipid type, we used *a*_*o*_ equal to the partial area per phosphate times the number of phosphate headgroups per lipid of that type (i.e., times 1 for POPC/DOPE and times 2 for CL) and *a*_*h*_ equal to the partial area per lipid acyl chain of times the total number of acyl chains per lipid of that type (i.e., times 2 for POPC/DOPE and times 4 for CL). This yields an estimate of the average value of *P*, quantifying the effective shape of the lipids within the bilayer.

#### 2.3.9 Hydrophobic thickness

The hydrophobic thickness, *h*_*C*_, of each bilayer was approximated by estimating average distance between the C2 carbon atoms of the lipid acyl chains (100, 101), computed using a 2-dimensional grid mapping tesselation procedure (see thickness calculations from Allen et al. (94) or Gapsys et al. (95)). This procedure was the same as that used in estimating the partial area per phosphate (Section 2.3.3), except the positions of the C2 carbon atoms of the lipid acyl chains projected into the xy-plane were used as the reference points for distance calculations and subsequent mapping of lipids to the grid elements, and in addition to mapping lipids to grid elements, the z-positions of the C2 atoms were also mapped to the grid elements. The hydrophobic distance for a snapshot was then computed as the average z-distance between corresponding grid elements of the opposing bilayer leaflets.

#### 2.3.10 P-P thickness

The phosphorous to phosphorous (P-P) thickness, *h*_*PP*_, of each bilayer was computed in the same manner as the hydrophobic thickness (Section 2.3.9), except that the positions of the phosphorous atoms of the lipid headgroup phosphates projected into the xy-plane were used as the reference points for distance calculations, and subsequent mapping of lipids and z-positions to the grid elements.

#### 2.3.11 Lipid lengths

The length of each individual lipid was estimated by computing the distance between the position of the lipid’s headgroup phosphorous atom and the center of mass of its terminal acyl chain carbons. The average lipid length of lipid type *l, L*_*l*_, was then estimated by taking the ensemble average of the lengths over lipids of type *l*.

#### 2.3.12 Bilayer surface fluctuations

We examined the bilayer surface fluctuations using the rectangular grid based tesselation procedure with grids generated in the same manner as described in Section 2.3.10 for the P-P thickness, but additionally, the area per grid element was fixed to 1 Å^2^ and the z-positions mapped to the grid elements were filtered through a Gaussian filter with *σ* = 5.0Å; the filter reduced noise and smoothed the surface representation generated by the grid assignments. For simplicity we only analyzed the upper leaflet of the bilayer. At each snapshot, a new grid was generated and the surface fluctuations or roughness was quantified by computing the standard deviation of the z-values for grid points that compose the surface, *σ*_*z*_. After analyzing all the snapshots from a trajectory, the time average, maximum, and minimum values of the *σ*_*z*_ values were computed.

#### 2.3.13 Correlation between local bilayer surface deflections and lipid molecule localization

The correlation between local bilayer surface deflections and lipid molecule localization was estimated using the procedure described in Koldsø et al. (102) (under the section “Correlation between bilayer surface curvature and the clustering of lipid molecules”). At each snapshot of the analysis the lateral area (xy box-dimensions) of the bilayer was divided into blocks; each leaflet is considered individually. Then the cross correlation between the deflection of lipids along the bilayer normal (in our case the z-dimension) relative to the mean position within the blocks and the local lipid composition within that block are estimated, yielding a normalized correlation for each lipid species:

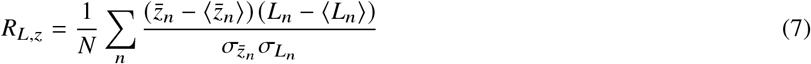

where *L*_*n*_ is the number of lipids of a given species in a grid box *n*, 〈*L*_*n*_〉 is the average of *L*_*n*_ over all grid boxes, 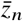 is the average *z* coordinate *z*_*n*_ of the lipid reference atom(s) (excluding the current lipid species being calculated) within grid block *n*, and 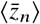 is the average of 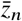 across all grid boxes. 3×3 grids (~48 × 48 Å^2^ grid elements) were used for the computation. The correlations for each leaflet at each snapshot of the analysis were time averaged to yield estimates of the correlation between bilayer surface curvature and the clustering of lipid molecules at each leaflet. The correlation for the upper and lower leaflet values of each lipid type were then averaged according to

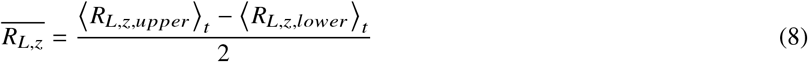

to yield a composite estimate combining the upper and lower leaflet values.

#### 2.3.14 Area compressibility modulus

The isothermal area compressibility modulus was estimated using the lateral area variance formula (103),

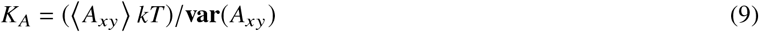

where *A* is the area in the lateral dimensions of the bilayer, *k* is the Boltzmann constant, and *T* the absolute temperature; the angle brackets denote the time average and **var()** the variance. The lateral bilayer area was approximated by the product of the x and y simulation box-dimensions. Measurements of the lateral box area were collected from the simulation trajectory at 2 ns intervals. The area measurements were then used with Equation 9 to yield final estimates of the area compressibility modulus.

#### 2.3.15 Bending modulus

The bending (or curvature) modulus of the bilayers, *K*_*C*_, was estimated from *K*_*A*_ and *h*_*C*_ via a polymer brush model (104, 105),

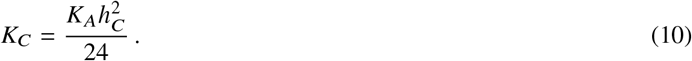

#### 2.3.16 Diffusion coefficients

The diffusion coefficients were estimated by applying the mean square displacement relation of the Einstein model of Brownian motion,

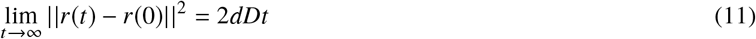

where the left hand side is the mean squared displacement as a function of time *t*, and on the right hand side *d* is the dimensionality, and *D* the diffusion coefficient. The mean squared displacement of each lipid type (averaged over all lipids of that type) was computed for the lateral motion of the center of mass of lipids in each leaflet of the bilayer; the lipid coordinates were adjusted at each step to remove the center of mass motion of the bilayer before estimating the mean squared displacements. The mean squared displacement curves were then fitted with a linear equation, the slope of which was used to estimate the diffusion coefficient according to Equation 11; note that *d* = 2 for 2-dimensional lateral motion.

## 3 RESULTS

### 3.1 Cardiolipin does not alter the bilayer lipid phase of model mitochodrial membranes

We first considered whether CL can modulate the phase behavior and packing of membrane lipids, which can in turn affect protein-membrane interactions (106), the protein-ligand interactions of embedded proteins (107), and membrane permeability (108). To characterize the bilayer lipid phase of the model mitochondrial membranes, we computed the average hexagonal packing order parameter for each bilayer as described in Section 2.3.1. This order parameter is unity for a perfect hexagonal packing and zero in the absence of hexagonal packing. All the model mitochondrial membranes had an order parameter value of 0.17 which is consistent with a liquid disordered phase (93), and indicated that CL content up to 15 mol% had no effect on the bilayer lipid phase in this geometry.

### 3.2 Cardiolipin increases the average area per lipid molecule in ternary mixtures with PC and PE lipids

Many of the membrane structural parameters are directly affected by the lateral packing density of the membrane lipid molecules. To characterize the lateral packing density of the lipid molecules in our model mitochondrial membranes we estimated the average area per lipid molecule as described in Section 2.3.2. The values obtained from our simulations are reported in Table 1, along with other previously reported simulation estimates for the average area per lipid molecule in similar planar ternary lipid bilayers from the literature. The data shows that increasing the mole percentage of CL increases the average area per lipid molecule. Our simulation data further indicates that the increase is non-linearly relative to the increase in CL content within the simulated concentration range. The increase of area per lipid molecule with CL content is consistent with trends reported for monolayer experiments (42–45, 109) (at given surface pressure) and bilayer simulations (49–51) of CL in binary mixtures with PC or PE lipids. Furthermore, since the total number of lipids was fixed in our simulations, the increase in the average area per lipid indicates that the average lateral area of the bilayer is expanded by the replacement of POPC and DOPE lipid molecules with CL lipid molecules. This observation fits intuitively with the fact that CL molecules are much larger than there two-acyl-chain counterparts and have average area per lipid molecule values exceeding 100 Å^2^ within pure CL bilayers in the liquid phase (105, 110).

**Table 1:**
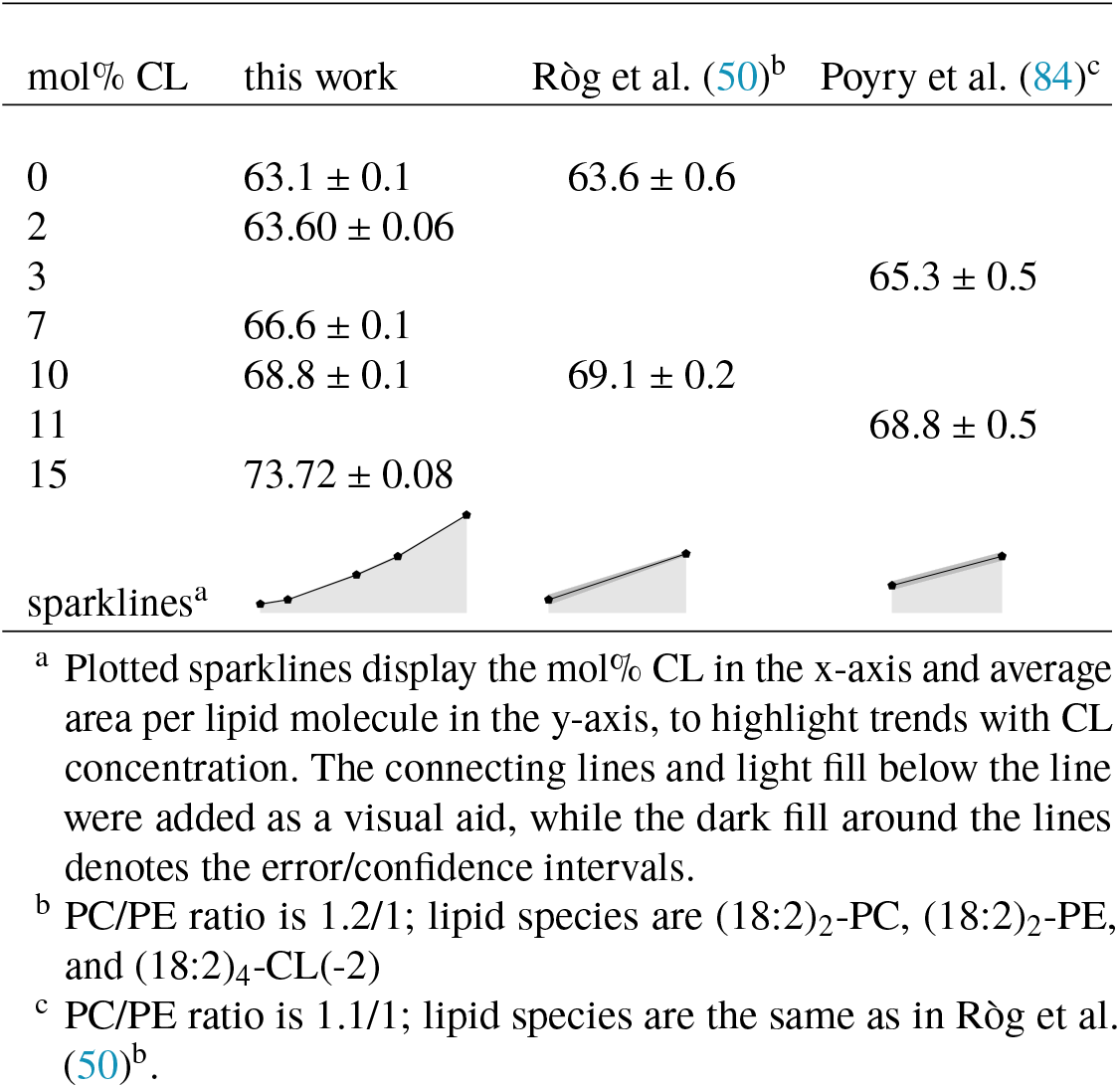
Average area per lipid molecule at various concentrations of CL in mixtures of PC and PE lipids.

### 3.3. Ternary mixtures of cardiolipin, PC and PE lipids with cardiolipin less than 15 mol% have slightly higher lateral headgroup and acyl chain packing densities as compared to corresponding cardiolipin free binary PC and PE lipid bilayer systems

The average area per lipid did not take into account the unique diglycerophospholipid structure of CL, which is essentially equivalent to two glycerophospholipids. Therefore, we further characterized the lateral lipid packing density by calculating both the average area per lipid phosphate group and the average area per lipid acyl chain, as well as their corresponding partial molar quantities.

The average area per lipid phosphate headgroup (abbreviated hereafter as average area per phosphate) was computed in the manner described in Section 2.3.3. The values obtained from our simulations are reported in Table 2, together with corresponding literature results for the average area per phosphate. It can be seen from our simulation data that the addition of up to 10 mol% CL into a mixture of PC and PE lipids results in a relative decrease in the average area per phosphate as compared to the CL free bilayer, with essentially no difference between the results for the 2, 7, and 10 mol% systems. The data from Ròg et al. (50) indicate that the addition of 10 mol% CL decreases the average area per phosphate as compared to the CL free bilayer, while the data from Poyry et al. (84) indicate that increasing the CL content from 3 to 11 mol% also results in a relative decrease in the average area per phosphate. Altogether, these results indicate that at low concentrations CL tends to decrease the average area per phosphate in ternary mixtures with PC and PE lipids; the relative differences in the levels of area per phosphate as shown by our simulation data and those of Ròg et al. (50) and Poyry et al. (84) may be attributed, at least in part, to differences in length and saturation of the PC and PE lipid acyl chains between our studies, as well as differences in the relative proportion of PC to PE lipids. In contrast to the relative decreases in area per phosphate at lower concentrations, our simulation data indicated a relative increase in area per phosphate at 15 mol% CL as compared to both the other CL containing bilayers and the CL free bilayer that we modeled. The average area per phosphate of our model mitochondrial membranes therefore exhibited a non-monotonic dependence on CL content.

**Table 2:**
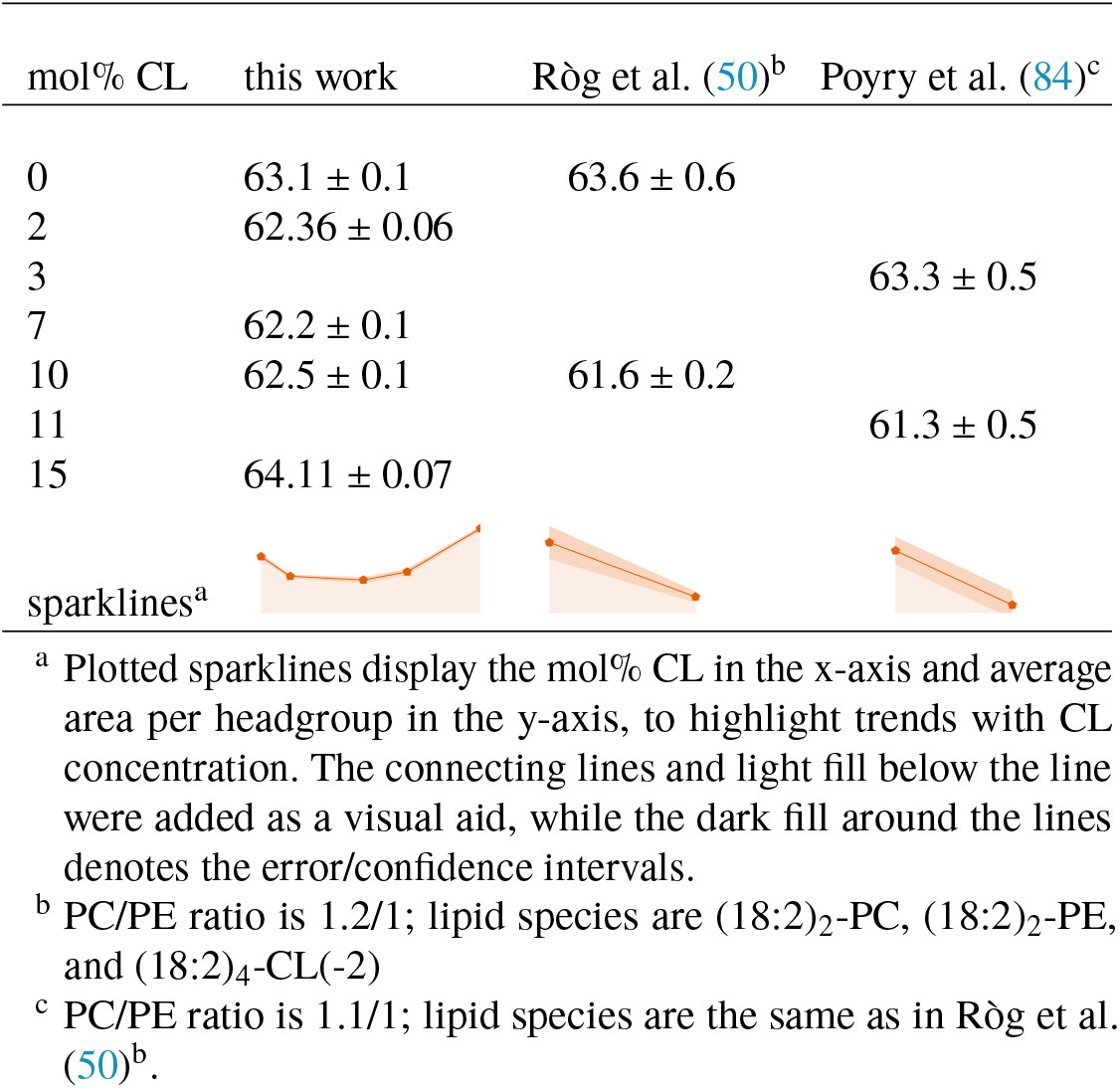
Average area per lipid phosphate headgroup; recall that each CL molecule has two phosphate headgroups. The average area per lipid phosphate headgroup was obtained from Ròg et al. (50) by multiplying the quantity reported therein as average area per hydrocarbon chain by a factor of 2. The average area per lipid phosphate headgroup was obtained from Poyry et al. (84) as the average area per lipid quantity reported therein.

| mol% CL | this work | Røg et al. (50)<sup>b</sup> | Poyry et al. (84)<sup>c</sup> |
| --- | --- | --- | --- |
| 0 | 63.1 ± 0.1 | 63.6 ± 0.6 |  |
| 2 | 62.36 ± 0.06 |  |  |
| 3 |  |  | 63.3 ± 0.5 |
| 7 | 62.2 ± 0.1 |  |  |
| 10 | 62.5 ± 0.1 | 61.6 ± 0.2 |  |
| 11 |  |  | 61.3 ± 0.5 |
| 15 | 64.11 ± 0.07 |  |  |
| sparklines<sup>a</sup> |  |  |  |

<sup>a</sup> Plotted sparklines display the mol% CL in the x-axis and average area per headgroup in the y-axis, to highlight trends with CL concentration. The connecting lines and light fill below the line were added as a visual aid, while the dark fill around the lines denotes the error/confidence intervals.
<sup>b</sup> PC/PE ratio is 1.2/1; lipid species are (18:2)<sub>2</sub>-PC, (18:2)<sub>2</sub>-PE, and (18:2)<sub>4</sub>-CL(-2)
<sup>c</sup> PC/PE ratio is 1.1/1; lipid species are the same as in Røg et al. (50)<sup>b</sup>.

To better understand the effects that CL has on the headgroup packing of the different lipid types, we estimated the partial area per phosphate of each lipid type using the approach described in Section 2.3.4. The results of this analysis are reported in Table 3. Our simulation data indicated that the partial area per phosphate of POPC and DOPE lipid molecules only underwent a small decrease when 2 mol% CL was added to the bilayer. Further increasing the mol% of CL resulted in small increases in their partial areas per phosphate. Interestingly, the partial areas per phosphate of POPC and DOPE lipids at 10 mol% either match or exceed their values in the CL free bilayer, suggesting that the relative decrease in the average area per phosphate maintained at 10 mol% CL (see Table 2) is primarily driven by the lower partial area per phosphate of the CL molecules and that the headgroup area of the PC and PE lipids are no longer compressed by the inclusion of CL. The partial area per phosphate of CL molecules increased monotonically but non-linearly as their concentration increased. In all cases, the 15 mol% CL system exhibited the largest differences from the other systems. Finally, as shown by our data, the area occupied by the headgroups of each lipid type followed the general trend POPC>DOPE>CL.

**Table 3:**
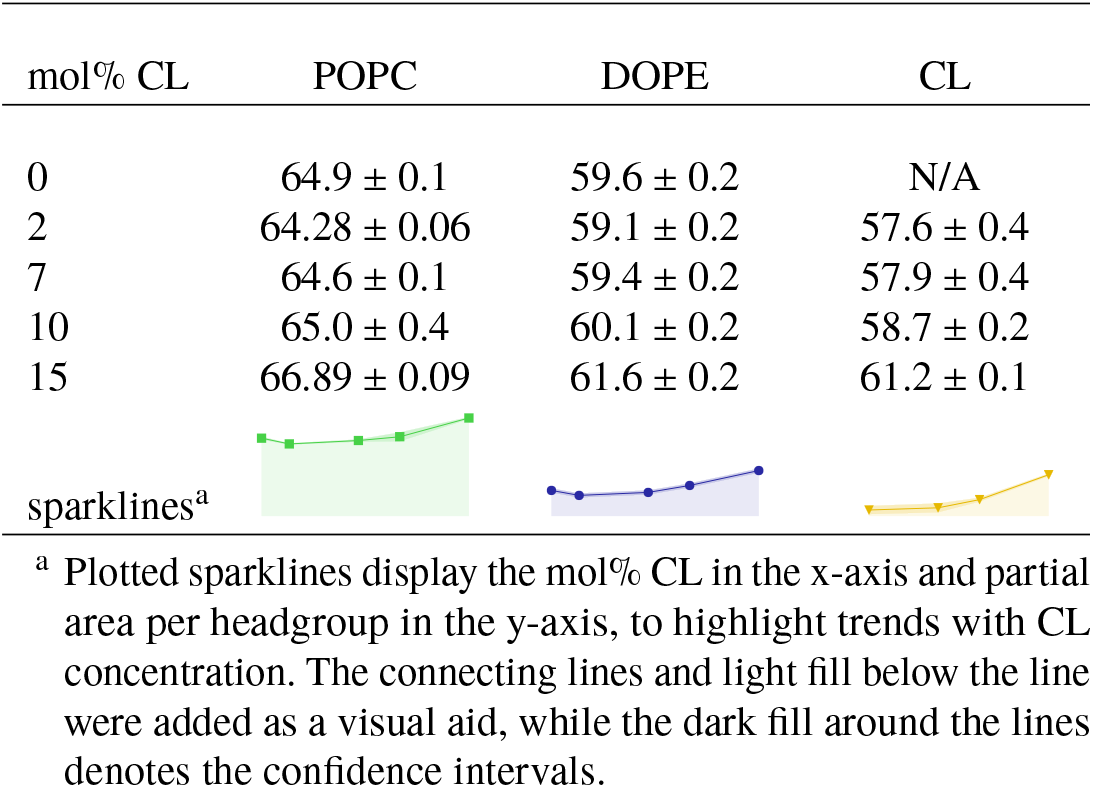
Partial areas per phosphate for each lipid type.

| mol% CL | POPC | DOPE | CL |
| --- | --- | --- | --- |
| 0 | 64.9 ± 0.1 | 59.6 ± 0.2 | N/A |
| 2 | 64.28 ± 0.06 | 59.1 ± 0.2 | 57.6 ± 0.4 |
| 7 | 64.6 ± 0.1 | 59.4 ± 0.2 | 57.9 ± 0.4 |
| 10 | 65.0 ± 0.4 | 60.1 ± 0.2 | 58.7 ± 0.2 |
| 15 | 66.89 ± 0.09 | 61.6 ± 0.2 | 61.2 ± 0.1 |
| sparklines<sup>a</sup> |  |  |  |

<sup>a</sup> Plotted sparklines display the mol% CL in the x-axis and partial area per headgroup in the y-axis, to highlight trends with CL concentration. The connecting lines and light fill below the line were added as a visual aid, while the dark fill around the lines denotes the confidence intervals.

In addition to examining the packing densities of the lipid headgroups, we also computed the average and partial areas per lipid acyl chain. The average area per lipid acyl chain was estimated in the manner described in Section 2.3.5 and the partial areas per lipid acyl chain of each lipid type were computed using the method outlined in Section 2.3.6. The results of these analyses are reported in Table 4. As can be seen from in our data, the trends in acyl chain areas are similar to those seen for the headgroups. The average and partial areas for POPC and DOPE were non-monotonic with CL concentration, showing a slight compression of the areas for 2-10 mol% CL relative to the CL free bilayer along with a relative expansion at 15 mol% CL. Since the average area per lipid acyl chain is related to the average area per phosphate by a constant multiplicative factor, it has the same trends as that quantity. However, the partial areas per lipid acyl chain for POPC and DOPE were not perfectly correlated with their corresponding partial areas per phosphate, remaining compressed at 10 mol% CL relative to the CL free system; their trends are much more closely aligned with the average area per lipid acyl chain. Additionally, the partial areas per lipid acyl chain are much more consistent across the different lipid types than the corresponding headgroup areas. With the headgroup areas there was clear trend in relative sizes. However, with the acyl chain areas the values for POPC and DOPE are indistinguishable, while CL has only a slight tendency to larger areas. Presumably, the higher density of the lipid acyl chains within the bilayer forces a more uniform packing of the lipid tails.

**Table 4:**
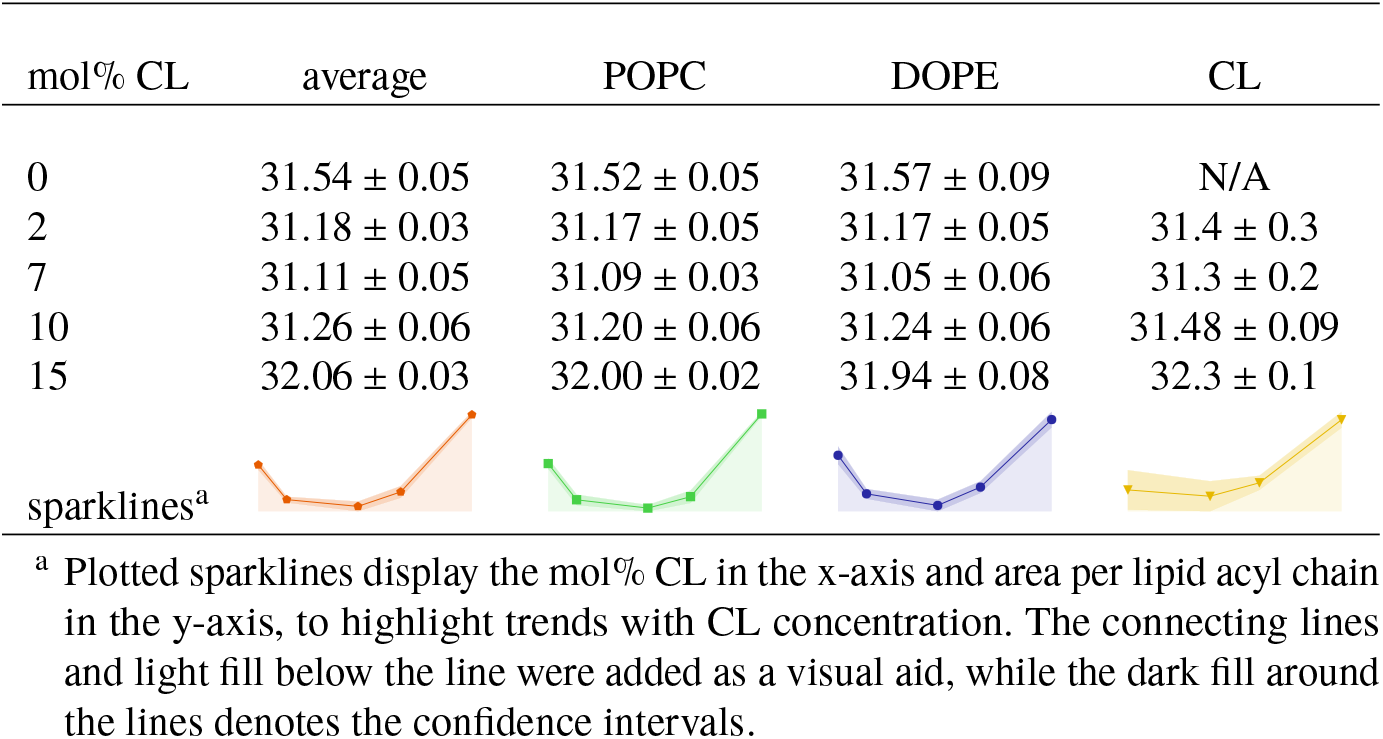
Average and partial areas per lipid acyl chain. Note that POPC and DOPE each have two acyl chains, while CL has four.

| mol% CL | average | POPC | DOPE | CL |
| --- | --- | --- | --- | --- |
| 0 | 31.54 ± 0.05 | 31.52 ± 0.05 | 31.57 ± 0.09 | N/A |
| 2 | 31.18 ± 0.03 | 31.17 ± 0.05 | 31.17 ± 0.05 | 31.4 ± 0.3 |
| 7 | 31.11 ± 0.05 | 31.09 ± 0.03 | 31.05 ± 0.06 | 31.3 ± 0.2 |
| 10 | 31.26 ± 0.06 | 31.20 ± 0.06 | 31.24 ± 0.06 | 31.48 ± 0.09 |
| 15 | 32.06 ± 0.03 | 32.00 ± 0.02 | 31.94 ± 0.08 | 32.3 ± 0.1 |
| sparklines<sup>a</sup> |  |  |  |  |

<sup>a</sup> Plotted sparklines display the mol% CL in the x-axis and area per lipid acyl chain in the y-axis, to highlight trends with CL concentration. The connecting lines and light fill below the line were added as a visual aid, while the dark fill around the lines denotes the confidence intervals.

### 3.4 Cardiolipin molecules repel one another, interacting preferentially with POPC molecules

Although CL lipid molecules are miscible in binary mixtures with PC lipid molecules(42, 43, 111), formation of laterally segregated domains has been reported for binary mixtures of CL and PE lipids (40, 43, 111) and in a ternary mixture of PC, PE, and CL (112). Additionally, CL enriched sub-domains within the outer mitochondrial membrane have been suggested to serve as an activating platform for mitochondrial apoptosis (5). To explore lipid aggregation, we characterized the lateral organization and mixing behavior of our model mitochondrial membranes. To do this, we characterized the lipid-lipid interactions and their lateral organization within the mitochondrial bilayers using a nearest-neighbour analysis and estimated the relative fractions of lipids interacting with their nearest neighbors as described in Section 2.3.7. The results of this analysis are displayed in Figure 3.

**Figure 3:**
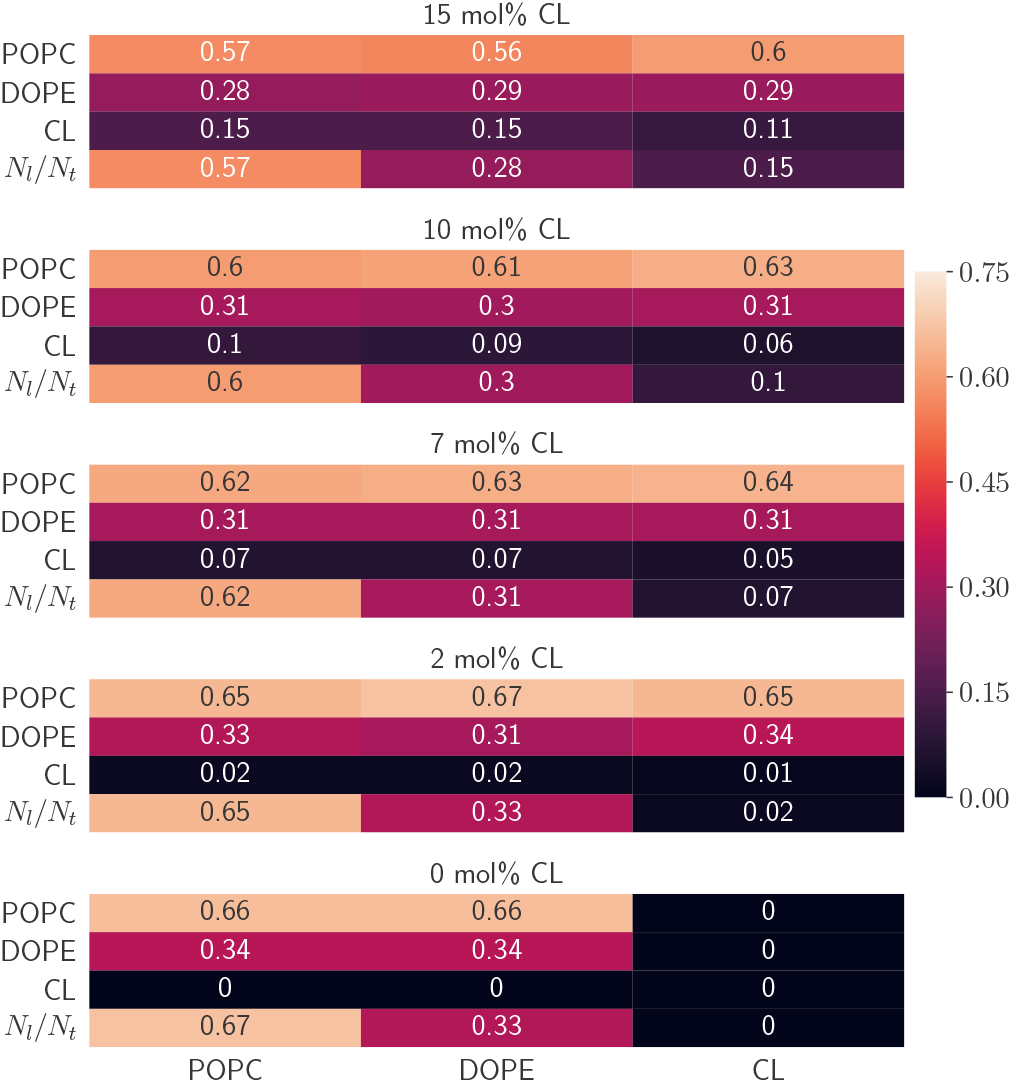
Lipid-lipid fractional interactions from nearest neighbor analysis for each lipid type and model system: computed as described in Section 2.3.7. The rows labeled *N*_*l*_/*N*_*t*_ designate the mole fractions of each lipid which are provided for reference. Fraction computations are centered on lipids along the x-axis. Errors are less than 0.02.

In general, the CL-CL interaction fractions are consistently lower than the CL’s mole fraction in each system as can be seen from the data in Figure 3. This suggests that the CL-CL interactions are less favourable, presumably due to repulsion between the negatively charged headgroups. However, it is possible that such a result could be an artifact of CL lipid molecules low proportion and slow diffusion within the bilayers. Therefore, we also analyzed the time-course of the CL-CL interaction fractions for each CL-containing system. The results of this analysis are displayed in Figure 4. As can be seen in Figure 4, While the fluctuations present in the 2 and 7 mol% CL systems obscure the small difference between the average value of *f*_CL-CL_ and CL’s mole fraction, the time-course data for the interactions in the 10 and 15 mol% CL clearly show that despite oscillations the CL-CL fractional interactions tend to be lower than the mole fraction of CL in those systems. These results are consistent with those displayed in Figure 3, and they further confirm that, at least in the timescale of these simulations, the CL-CL interactions appear to be repulsive.

**Figure 4:**
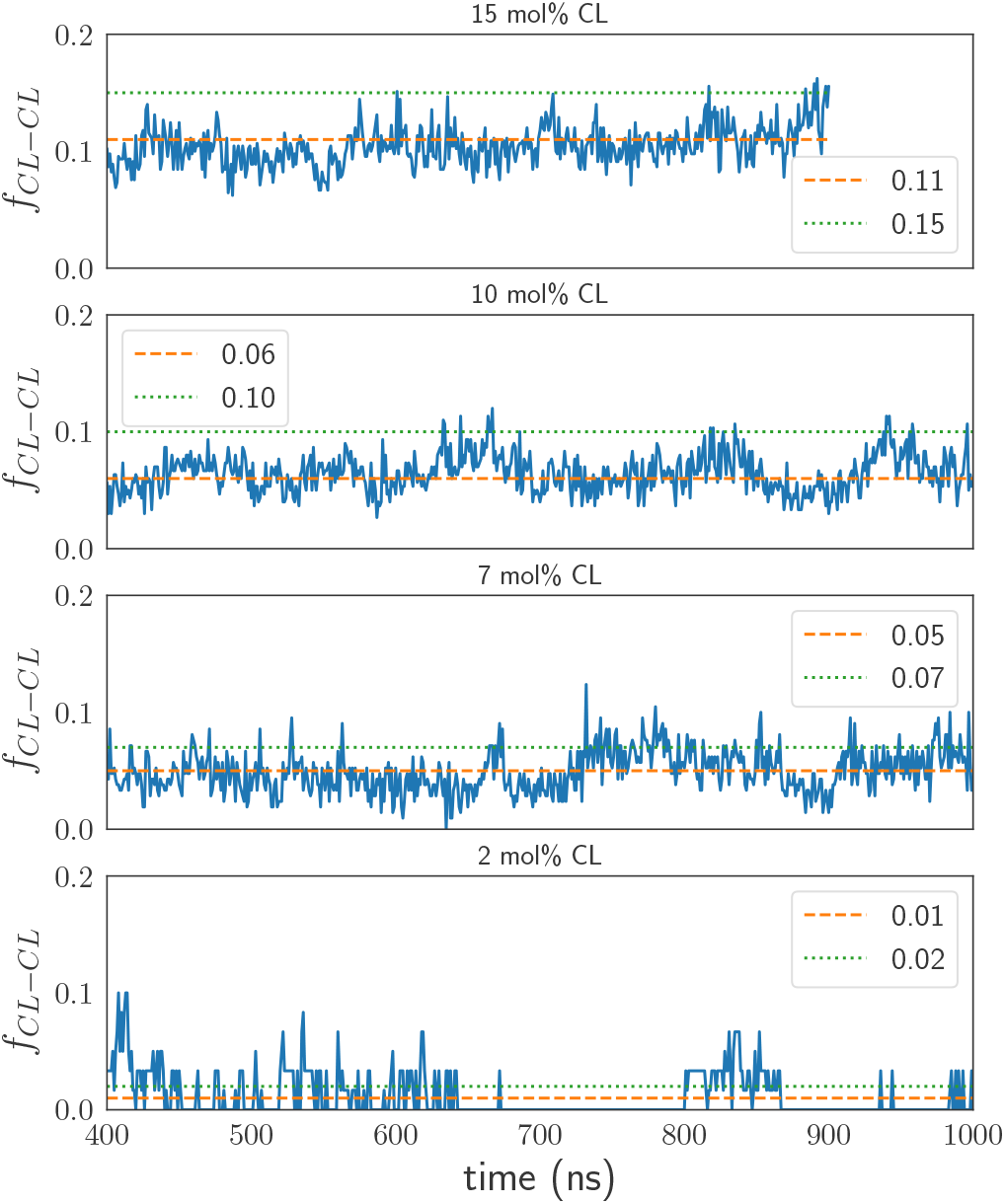
CL-CL fractional interactions (*f*_CL-CL_) from nearest neighbor analysis plotted versus time.

Additionally, the CL-POPC interaction fractions exceeded POPC’s mole fraction in the 7, 10 and 15 mol% CL systems, suggesting that the CL molecules interact preferentially with POPC molecules. CL molecules have been reported to have a stronger condensing effect in binary mixtures with PC lipids than in binary mixtures with PE lipids (50, 51), suggesting that CL molecules tend to have stronger attractive interactions with PC lipids than with PE lipids. Ròg et al. (50) also reported that PC molecules preferentially formed charge pairs with CL molecules within a bilayer composed of a ternary mixture of PC, PE, and CL lipid molecules.

### 3.5 CL-dependent changes in lipid packing do not induce significant changes in bilayer thickness

Lipid monolayers and bilayers exhibit many of the properties of an elastic sheet (113, 114). Under this model, alterations in the bilayer area result in commensurate alterations in the bilayer thickness. Since the changes in CL content resulted in minor changes to the lipid packing areas, we estimated some of the bilayer thickness properties to see if they also changed with CL content. The results of these analyses are reported in Table 5. First, we estimated the hydrophobic thickness, *h*_*C*_, of each bilayer using the method described in Section 2.3.9. The hydrophobic thickness quantifies the thickness of a bilayer’s hydrophobic core (hydrocarbon region) and is particularly important for protein-membrane interactions, driving integral protein-membrane interaction energetics through hydrophobic mismatch (64, 115). As the data show, the variation of *h*_*C*_ with CL content was minimal (< 1Å), but the changes followed the trends (negatively correlated) in the areas per phosphate and acyl chain (see Section 3.3). The *h*_*C*_ values, 28 – 29Å, agree with the average length (28.6 ± 1.4 Å) of the hydrophobic transmembrane domains of IMM proteins reported by Pogozheva et al. (116).

**Table 5:**
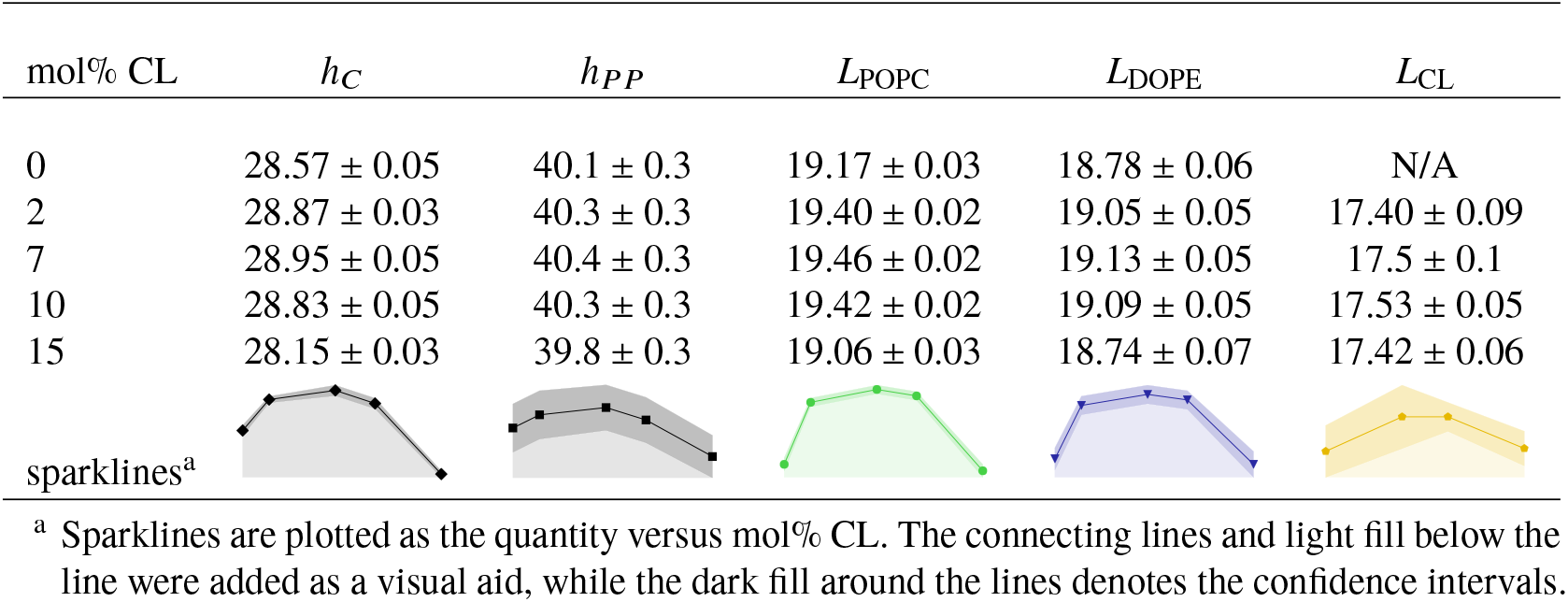
Hydrophobic thickness (*h*_*C*_), P-P thickness (*h*_*PP*_), and lipid lengths (*L*_*l*_). Results are reported in units of Å.

| mol% CL | $h_C$ | $h_{PP}$ | $L_{POPC}$ | $L_{DOPE}$ | $L_{CL}$ |
| --- | --- | --- | --- | --- | --- |
| 0 | $28.57 \pm 0.05$ | $40.1 \pm 0.3$ | $19.17 \pm 0.03$ | $18.78 \pm 0.06$ | N/A |
| 2 | $28.87 \pm 0.03$ | $40.3 \pm 0.3$ | $19.40 \pm 0.02$ | $19.05 \pm 0.05$ | $17.40 \pm 0.09$ |
| 7 | $28.95 \pm 0.05$ | $40.4 \pm 0.3$ | $19.46 \pm 0.02$ | $19.13 \pm 0.05$ | $17.5 \pm 0.1$ |
| 10 | $28.83 \pm 0.05$ | $40.3 \pm 0.3$ | $19.42 \pm 0.02$ | $19.09 \pm 0.05$ | $17.53 \pm 0.05$ |
| 15 | $28.15 \pm 0.03$ | $39.8 \pm 0.3$ | $19.06 \pm 0.03$ | $18.74 \pm 0.07$ | $17.42 \pm 0.06$ |
| sparklines<sup>a</sup> |  |  |  |  |  |

<sup>a</sup> Sparklines are plotted as the quantity versus mol% CL. The connecting lines and light fill below the line were added as a visual aid, while the dark fill around the lines denotes the confidence intervals.

In general, membrane thickness plays an important role in membrane protein-membrane interactions (Li2012, Karabadzhak2018) (117), affecting the structure (53, 64, 116, 118), dynamics (119, 120) and distribution (62) of integral and associated proteins, as well as directly affecting membrane permeability (121). To gauge the overall thickness of the bilayer membranes, we computed the average phosphorous to phosphorous (P-P) thickness, *h*_*PP*_, of the bilayers using approach described in Section 2.3.10. As the data in Table 5 show, the *h*_*PP*_ values appear to follow a similar trend with CL content as the *h*_*C*_ values, but the small variations in *h*_*PP*_ are obscured by statistical error. Regardless, values of *h*_*PP*_ ≈ 40Å compare favorably with average P-P thickness (42Å) reported for the bilayer with a ternary mixture of PC/PE/CL simulated by Ròg et al. (50), as well as the average distance between phosphate groups (41 ± 1Å reported by Dahlberg and Maliniak (49) for coarse grain simulation of a binary mixture of teraoleoyl CL (TOCL, 9.2%) in POPC.

To better understand the molecular contributions of the different lipid types to the bilayer thickness, we estimated the average length of lipids, *L*_*l*_, using the approach described in Section 2.3.11. As can be seen from the data in Table 5, similar to the membrane thickness metrics the variation of the *L*_*l*_ values with CL content were minimal for POPC and DOPE (< 0.5 Å), but followed the same overall trends; the changes in *L*_*l*_ with CL content correlated reasonably well to trends those observed for the partial areas per lipid acyl chain (see Section 3.3). Within the simulated concentration range, the average length of CL molecules was virtually unaffected by their concentration. Additionally, the differences between lipid types corresponded well to acyl chain unsaturation, following the trend *L*_*POPC*_ > *L*_*DOPE*_ > *L*_*CL*_.

### 3.6 Increasing Cardiolipin concentration decreases bilayer elasticity

A membrane’s elastic properties characterize how it responds to deformations, such as stretching and bending. Membrane elastic properties affect the energetics of protein-membrane interactions and thereby influence integral membrane function (59, 64, 122, 123). To characterize the elasticity of the model mitochondrial bilayers we estimated the two principal bilayer elastic moduli (113, 114): the area compressibility (or area expansion) modulus and the bending (or curvature) modulus.

The area compressibility modulus, *K*_*A*_, characterizes the force required for lateral deformations of the bilayer surface area, while the bending modulus, *K*_*C*_ (sometimes denoted as *K*_*B*_), characterizes the energetic cost of curvature inducing bending deformations along the bilayer normal. Estimates of *K*_*A*_ and *K*_*B*_ for each bilayer were computed using the methods described in Section 2.3.14 and Section 2.3.15, respectively. Our simulation results from these analyses are reported in Table 6. As the data show, both *K*_*A*_ and *K*_*C*_ of the model mitochondrial membranes were increased by the inclusion of CL into the bilayers, suggesting that CL incorporation decreased the elasticity of the membranes.

**Table 6:**
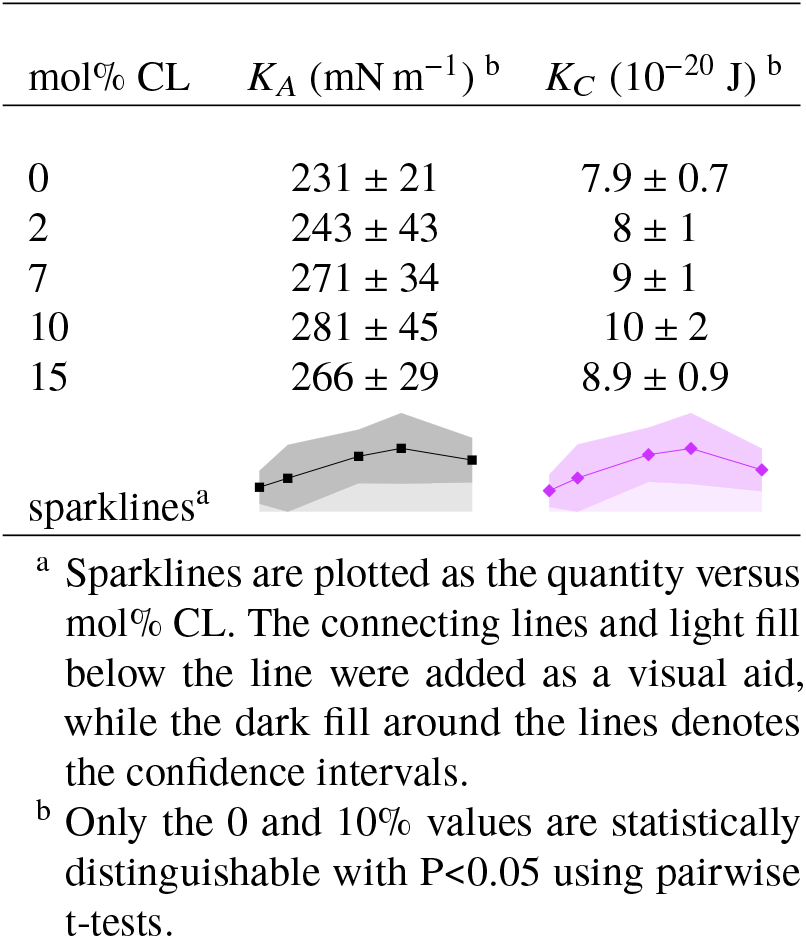
Mechanical moduli. Errors are reported as 1.96 times the standard error of the mean (i.e., 1.96 × SEM) and represent 95% confidence intervals.

| mol% CL | $K_A$ (mN m<sup>-1</sup>)<sup>b</sup> | $K_C$ (10<sup>-20</sup> J)<sup>b</sup> |
| --- | --- | --- |
| 0 | 231 ± 21 | 7.9 ± 0.7 |
| 2 | 243 ± 43 | 8 ± 1 |
| 7 | 271 ± 34 | 9 ± 1 |
| 10 | 281 ± 45 | 10 ± 2 |
| 15 | 266 ± 29 | 8.9 ± 0.9 |
| sparklines<sup>a</sup> |  |  |

<sup>a</sup> Sparklines are plotted as the quantity versus mol% CL. The connecting lines and light fill below the line were added as a visual aid, while the dark fill around the lines denotes the confidence intervals.
<sup>b</sup> Only the 0 and 10% values are statistically distinguishable with $P < 0.05$ using pairwise t-tests.

### 3.7 Cardiolipin concentration in the membrane has a significant impact on lipid diffusion coefficients in the membrane

Fluidity (i.e., viscosity) is another physical property of membranes that is important for lipid distribution and protein-membrane interactions (55, 61, 62). Membrane fluidity plays a key role in controlling the diffusion (120, 124) and conformational changes of membrane bound proteins (53, 64), making the property particularly important in signal transduction cascades that depend on membrane bound diffusion-limited protein-protein interactions and protein conformational changes, such as in mitochondrial apoptosis regulation.

Membrane fluidity (or viscosity) is intrinsically linked to the lateral diffusion coefficients of its lipid constituents. Therefore, we estimated the lateral diffusion coefficients, *D*_*l*_, of each lipid species using the method described in Section 2.3.16. Our simulation results are reported in Table 7.

**Table 7:**
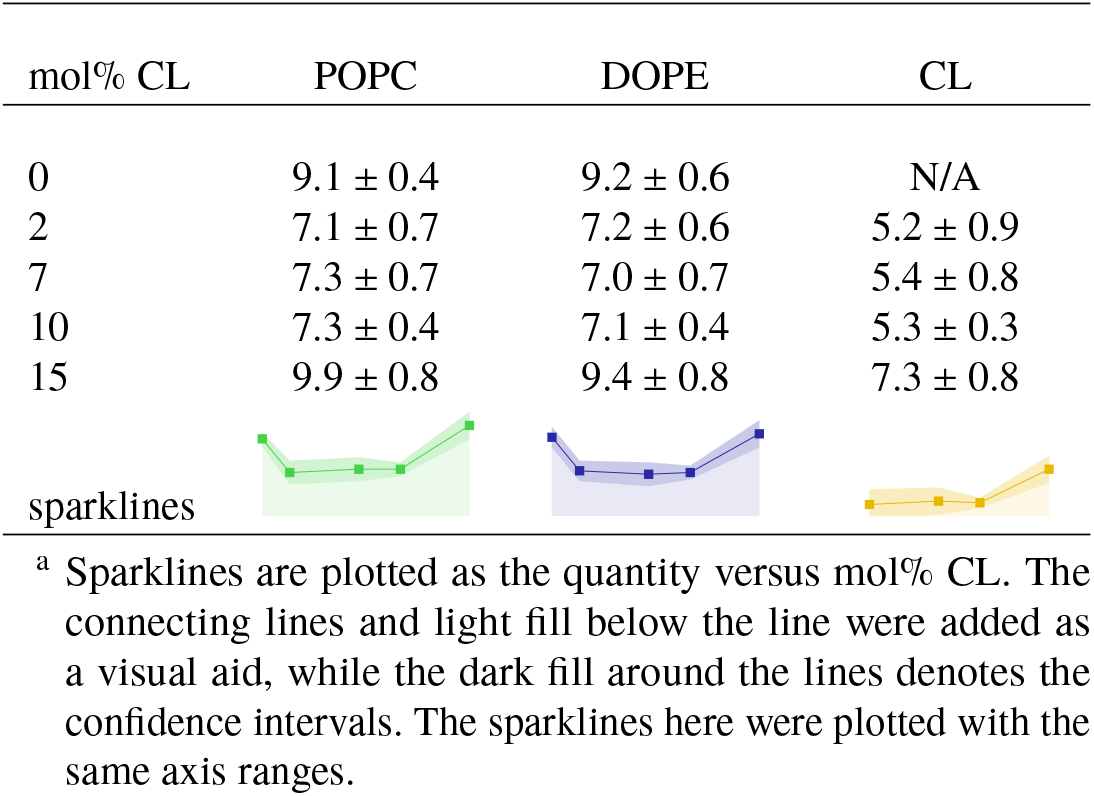
Lateral lipid diffusion coefficients, *D*_*l*_. Results are reported in units of 10^-8^ cm^2^/s.

| mol% CL | POPC | DOPE | CL |
| --- | --- | --- | --- |
| 0 | 9.1 ± 0.4 | 9.2 ± 0.6 | N/A |
| 2 | 7.1 ± 0.7 | 7.2 ± 0.6 | 5.2 ± 0.9 |
| 7 | 7.3 ± 0.7 | 7.0 ± 0.7 | 5.4 ± 0.8 |
| 10 | 7.3 ± 0.4 | 7.1 ± 0.4 | 5.3 ± 0.3 |
| 15 | 9.9 ± 0.8 | 9.4 ± 0.8 | 7.3 ± 0.8 |
| sparklines |  |  |  |

<sup>a</sup> Sparklines are plotted as the quantity versus mol% CL. The connecting lines and light fill below the line were added as a visual aid, while the dark fill around the lines denotes the confidence intervals. The sparklines here were plotted with the same axis ranges.

In general, the diffusion coefficients for POPC and DOPE match reasonable well with those expected from phospholipids in the liquid state (∼ 3 – 11 × 10^-8^ *cm*^2^/*s*) (125). The values for CL compare favorably to previously published diffusion coefficients for tetramyristol CL (∼5 × 10^-8^ *cm*^2^/*s*) (126) and tetraoleoyl CL (∼4 × 10^-8^ *cm*^2^ *s*) (127) in liquid phase unary CL bilayers. The trends in *D*_*l*_ values with CL content are correlated with changes in the partial areas per lipid acyl chain, consistent with the correlation between the lateral lipid packing density and lipid diffusion reported by Javanainen et al. (92). The resultant decrease in POPC and DOPE diffusion at at 2, 7, and 10% CL suggests a relative decrease in membrane fluidity at those concentrations, which is reversed at 15 % CL. Interestingly, even though the partial areas per lipid acyl chain were only condensed by about 3% at 2, 7, and 10% CL, *D*_*l*_ values for POPC and DOPE lipids were decreased by about 20 % at those concentrations. Thus, even though the changes to the membrane structural properties are subtle, corresponding changes in the lipid lateral diffusion are significant.

### 3.8 Although the effective packing shape of lipid molecules is cylindrical, they have small biases towards truncated conical shapes

Under the conditions we simulated CL(−2) lipids would be expected to be stable in the lamellar phase, but under certain pH and salt conditions lamellar CL membranes transition to a, nonlamellar, inverted hexagonal phase (39, 40). The transition is driven by an effective decrease in the head-to-tail size ratio as the CL headgroups are protonated or sufficiently screened. Similarly, PE lipids are able to form inverted hexagonal phases (128), and inverted hexagonal phases have also been observed for mixtures of CL and PE lipids (40, 43, 111). The ability to form inverted hexagonal phases signifies an inherent bias towards an inverted conical packing shape. Therefore, we quantified the effective shape of lipid molecules within our model mitochondrial bilayers by estimating the average values of the dimensionless packing shape parameter (97) based on their lateral packing areas using the approach described in Section 2.3.8; this parameter is equal to 1 for cylinder shaped lipid molecules, <1 for truncated cone shaped lipid molecules, and >1 for inverted truncated cone shaped lipid molecules. The results of this analysis are reported in Table 8. As can be seen from the data in Table 8, all the lipids have packing shape parameters ∼ 1 which indicates that the lipids are packed with effectively cylindrical shapes. Note that values near ∼ 1 were to be expected since the lipid systems we modeled are planar bilayers (lamellar phase), and lamellar bilayer lipids have cylindrical shapes with packing parameters ~ 1 (97, 98). However, CL and DOPE have a small but consistent bias to values >1, in accordance with a slight preference to take on truncated cone shapes. In contrast, POPC has a slight bias to value <1, suggesting a slight preference to inverted truncated cone shapes. And since averaging the packing parameters across lipid types results in an average value of 1.00, the slight differences in packing shape between POPC and CL/DOPE appear to be complementary.

**Table 8:**
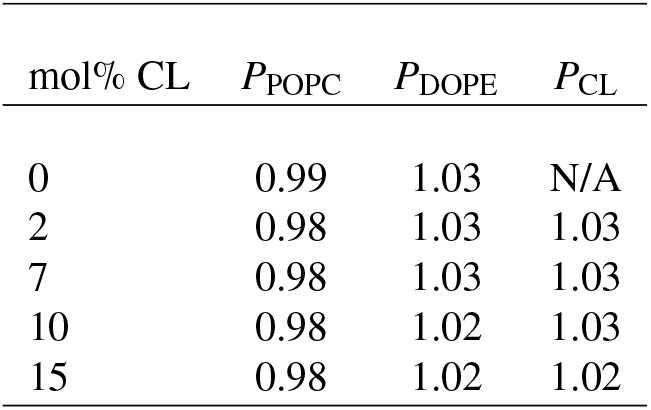
Effective critical packing parameters of each lipid type in the model mitochondrial membranes. Errors are 0.01 or less.

### 3.9 Cardiolipin and DOPE lipid molecules correlate to local negative deviations in the the bilayer surface curvature

Both CL and DOPE lipids have a propensity to form inverted hexagonal phases and are thus expected to have a preference towards inverted conical shapes and negative curvatures. However, as our estimates of the effective packing shape parameter (Section 3.8) indicated, all the lipids were constrained to approximately cylindrical shapes within the bilayer. Restriction of the lipids to less favorable packing shapes can result in curvature stress within the membrane monolayers (129) and a tendency towards spontaneous local curvature fluctuations Dahlberg and Maliniak (51). In the case of the interactions that regulate mitochondrial apoptosis, a curvature-dependent mechanism has been implicated in tBid-interactions (130, 131) while permeabilization via Bax-type molecules was reported to be sensitive to intrinsic monolayer curvature (132). Therefore, we evaluated the relationship between lipid localization and local curvature fluctuations within the bilayer surface. To quantify this relationship we computed the cross correlation between local bilayer surface deflections and lipid molecule localization, 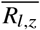, as described in Section 2.3.13. The results of this analysis are reported in Table 9. As the data show, DOPE and CL localization tended towards negative correlation coefficients, indicating a bias toward negative curvature deflections in the bilayer surface. Consistent with this result, Dahlberg and Maliniak (51) reported that CL induced negative spontaneous curvature in coarse grained lipid bilayer simulations. Additionally, Dahlberg and Maliniak (51) also reported that at low concentrations of a fully ionized CL, DOPE/CL bilayers had more negative spontaneous curvature than another system containing partially ionized CL in POPC.

**Table 9:** Cross correlations between local bilayer surface deflections and lipid molecule localization, 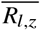.

| mol% CL | POPC | DOPE | CL |
| --- | --- | --- | --- |
| 0 | $0.0 \pm 0.1$ | $-0.3 \pm 0.1$ | N/A |
| 2 | $0.04 \pm 0.06$ | $-0.18 \pm 0.06$ | $-0.05 \pm 0.09$ |
| 7 | $0.02 \pm 0.08$ | $-0.2 \pm 0.1$ | $-0.2 \pm 0.1$ |
| 10 | $0.04 \pm 0.09$ | $-0.11 \pm 0.07$ | $-0.09 \pm 0.08$ |
| 15 | $0.1 \pm 0.1$ | $-0.3 \pm 0.1$ | $-0.1 \pm 0.1$ |

Our results show that the correlation coefficient for POPC lipids was zero in the 2-10 mol% systems. This is not too surprising because POPC is the highest mole fraction component in each of the bilayers, and therefore has the greatest weight in determining their average surface properties. Interestingly, POPC had a small positive correlation coefficient in the 15 mol% system, consistent with a its mild bias towards a truncated cone packing shape (Section 3.8).

### 3.10 The magnitude of bilayer surface fluctuations is not significantly affected by cardiolipin content

For a planar bilayer under zero tension, the surface is expected to be relatively flat. However, as can be seen in Figure 5, there were small deformations in the surface of the bilayers, contributing to roughness and small local changes in surface curvature. To determine if CL concentration affected the overall surface roughness and the magnitude of surface fluctuations we computed the standard deviation of the z-positions associated with the bilayer surface, including its time average, minimum, and maximum values from each simulation using the analysis methods described in Section 2.3.12. Our simulation results from this analysis are reported in Table 10. As the data indicate, increasing the mol% of CL did not have a significant effect on the overall surface roughness or the magnitude of surface fluctuations. We did not examine the low frequency bilayer undulations, however, previous coarse grained simulations (51) of binary PC/CL and PE/CL bilayers reported that increased CL content, particularly CL with a single −1 charge, resulted in greater deformability and larger undulations. Additionally, the results from another set of coarse grained simulations (133) suggested that CL increased a bilayer’s ability to deform when under strain.

**Table 10:** Standard deviation of the surface z-coordinates from the upper leaflet of the bilayers. Units are in Å.

| mol% CL | minimum | time average | maximum |
| --- | --- | --- | --- |
| 0 | 1.3 | 2.2 | 3.6 |
| 2 | 1.2 | 2.2 | 4.2 |
| 7 | 1.3 | 2.2 | 3.6 |
| 10 | 1.2 | 2.2 | 3.6 |
| 15 | 1.5 | 2.4 | 3.9 |

**Figure 5:**
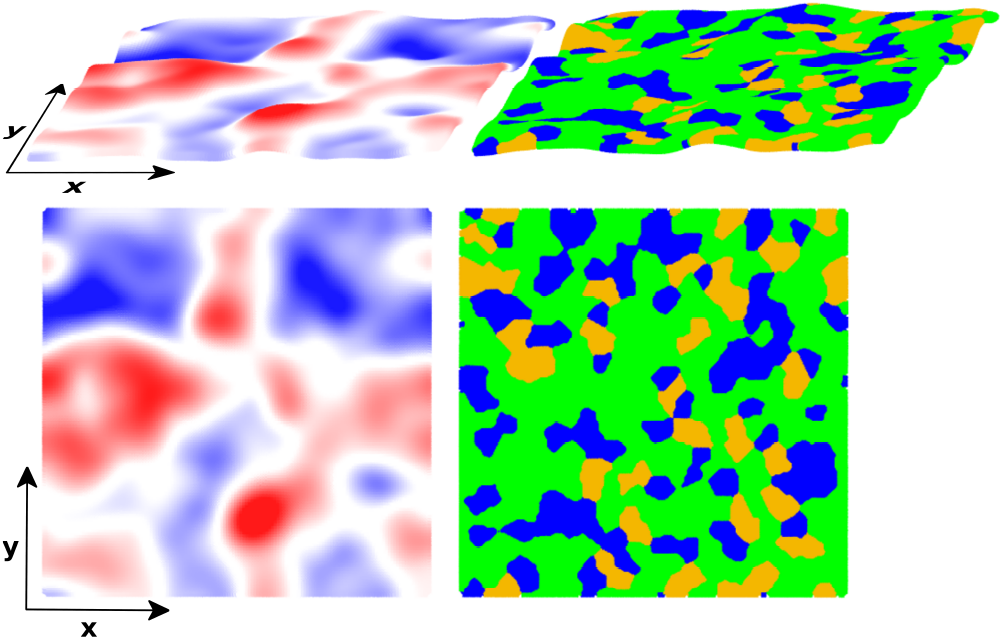
Sample snapshot taken from the 10 mol% CL system of the surface of the upper bilayer leaflet demonstrating surface roughness and curvature fluctuations. **Left:** heatmapped according to the ∆*z* value at each point of the surface; the heatmap is color coded blue → red: –4Å → 4Å. **Right:** color coded at point according to the lipid type; lipid colors are POPC:green, DOPE:blue, and CL:gold. The images were generated using the Tachyon renderer (http://jedi.ks.uiuc.edu/∼johns/raytracer/) in VMD (http://www.ks.uiuc.edu/Research/vmd/) (76, 77).

## 4 DISCUSSION

In this article, we have constructed five atomistic models of ternary lipid bilayer systems mimicking mitochondrial membranes. The membrane bilayers were square patches with sizes of ∼ (14-15 nm)^2^ and were composed of POPC, DOPE and varying proportions of cardiolipin (CL) molecules consistent with those in the outer (∼ 1 – 5 mol%, ∼ 15 mol% at contact sites) and inner (∼ 5 – 10 mol%) mitochondrial membranes. We characterized the properties of the membranes by microsecond molecular dynamics simulations. To understand how the CL molecules influenced the properties of these model mitochondrial, we estimated a variety of structural, mechanical, and dynamic bilayer properties from those simulations.

From our simulations, we found that CL content up to 15 mol% had only small effects on model membrane structural properties (see Section 3.1-Section 3.5). The addition of 2 mol% CL into the mixed POPC-DOPE matrix resulted in a slight bilayer condensation (Section 3.3) that was accompanied by a significant decrease in membrane fluidity (Section 3.7), as well as decreased membrane elasticity (Section 3.6). Further structural and fluidity changes with CL content were negligible between systems with 2 to 10 mol% CL, but elasticity was further decreased. Interestingly, the condensing effect was reversed for the 15 mol% system, which was slightly expanded relative to all other systems and exhibited greater fluidity; we suspect the transition could be driven by electrostatic repulsion between the CL molecules. The results from the simulations by Ròg et al. (50) indicated that the ternary mixture of PC, PE, and 10 mol% CL they modeled was only mildly condensed as compared to the corresponding CL free binary PC/PE bilayer (see Table 2). Similarily, the results from the simulations by Poyry et al. indicated a small condensation in a ternary lipid bilayer when going from 3 to 11 mol% CL (also Table 2). Additionally, fluorescence experiments performed by Khalifat et al. (21) suggested that while the addition of 10 % bovine cardiac CL to PC bilayers (at pH 7.4) lead to condensation, there was no discernible effect on mixed PC/PE bilayers. Taken together, these studies along with our data suggested that CL induces a mild condensing/ordering effect in mixed PC-PE lipid bilayers when the concentration of CL is < 15 mol%. Furthermore, our data indicated that the system with 15 mol% CL had greater mebrane fluidity. To the best of our knowledge, there are no other studies of similar ternary mixtures that examine the lipid packing properties at a CL concentration of ∼ 15 mol%. Therefore, we are unable to make a direct comparison for our results of the 15 mol% system.

Although the formation of laterally segregated domains has been reported for binary mixtures of CL and PE lipids (40, 43, 111) and for a ternary mixture of PC, PE, and 20 mol% CL (112), we did not find any evidence of domain formation within our model systems. Indeed, our analysis of the lateral lipid-lipid interactions and mixing suggested that lipids were for the most part well mixed (see Section 3.4). The one major exception were the CL-CL interactions which we found to be consistently lower than dictated by the CL mole fraction, suggesting that CL-CL interactions were at least somewhat repulsive on average. We suspect this effect is driven by electrostatic repulsion between CL molecules’ negatively charged headgroups. Regardless, it should be noted that the initial configurations of our systems were approximately well mixed, which may have biased the systems against domain formation within the timescales that we simulated. And although considerably larger than similar bilayer systems previously simulated, it is also possible the size of the bilayer patches used in our simulations were still too small to accurately gauge the formation of laterally segregated domains. Therefore, our result should be taken as suggesting an underlying repulsion between the negatively charged CL molecules but not definitive evidence against domain formation in these ternary PC, PE, and CL lipid systems.

Our analysis of the effective packing shape of the lipids revealed that the lipids took on near cylindrical shapes within the bilayers (see Section 3.8). And although analysis of the surface roughness indicated that CL content did not have a significant effect on the magnitude of surface roughness fluctuations (see Section 3.10), analysis of the correlation between CL localization and surface deflection revealed that CL had a tended to cause local deflections in the bilayer surface with negative curvature (see Section 3.9). A recent simulation study performed by Boyd et al. (133) reported that CL became locally enriched in the negative curvature regions of buckled bilayers, suggesting that lateral reorganization helped stabilize buckling deformations. Together with our results, these data suggest that CL is constrained to a non-favorable packing shape within bilayers that results in curvature frustration and a tendency to negative curvature fluctuations. As such, when the bilayer is perturbed by external forces into a highly curved state, as in the buckling simulations, CL responds by reorganizing into the negatively curved regions, alleviating its own inherent curvature strain and stabilizing the negatively curved regions of the bilayer.

## 5 CONCLUSIONS

In this work we have examined the cardiolipin-dependent properties in ternary lipid bilayer systems composed of PC, PE, and cardiolipin using atomistic molecular dynamics simulations. We modeled five lipid bilayers with cardiolipin content encompassing the natural range of cardiolipin content found in mitochondrial membranes, including the inner and outer mitochondrial membrane and their contact sites. From this series of simulations we estimated a variety of membrane properties, including structural quantities, mechanical moduli, and lipid diffusion. The properties we have characterized are known to influence the energy and kinetics of protein-membrane and protein-protein interactions at membranes (63, 64), and consequently affect membrane protein properties such as conformation (52–56), activity (57–60), and distribution (61, 62).

Our results indicated that CL content of 2-10 mol% induced a minor condensation of the bilayer lipids (∼1 – 3% reduction in the area occupied by the lipid acyl chains) that was accompanied by a decrease in membrane elasticity (∼ 20% increase in area compressibility modulus in the 10 % CL system) and lipid diffusion (∼ 20% decrease in the diffusion coefficients of both POPC and DOPE in all three systems). However, in contrast, the 15 mol% CL system underwent a small amount of bilayer expansion relative to the 2-10 mol% systems (∼ 2 – 3% increase in the area occupied by the lipid acyl chains) that was accompanied by a significant increase in the lipid diffusion (∼30 – 40% relative increase in the lipid diffusion coefficients). Overall, our analysis suggested that CL plays a minor role in altering membrane structural features in ternary lipid bilayers with a background matrix of PC and PE lipids, but can have significant effects on bilayer fluidity. Furthermore, our data indicated that these cardiolipin-dependent effects are non-monotonic, indicating a delicate balance between attractive and repulsive interactions within the bilayers.

Our model systems did not show any evidence of laterally segregated domain formation, but as noted in the Discussion, our simulations do not rule out such a phenomenon. Further analysis at larger length and longer time scales will likely be needed to further explore this particular phenomenon in ternary mixture models of mitochondrial membranes.

Finally, examination of the lipid packing shapes and curvature fluctuations suggested that CL has correlates to negative curvature deflections of the bilayer surface and likely induces negative curvature strain within the membrane monolayers.

This work contributes to a foundational understanding of the role of CL in altering the properties ternary lipid mixtures composed of PC, PE, and CL molecules. And since these are the primary components of mitochondrial membranes, this work also serves to illuminate the concentration-dependent role of CL in the mitochondrial membranes. However, our models did not capture the full complexity of the lipid milieu of mitochondrial membranes, including the inclusion of various minor components (e.g. PS, PI, and cholesterol) and the distributions of acyl chain length and saturation. Additionally, in our model membranes CL was distributed equally among bilayer leaflets, not accounting for the possibility of asymmetric lipid distribution. Additional experiment and simulation of more complex lipid mixtures will be required to determine what role these differences have on bilayer structure and dynamics.

## AUTHOR CONTRIBUTIONS

B.A.W., A.R., and C.F.L., designed the research. B.A.W. performed the research and analyzed the data. B.A.W., A.R., and C.F.L. wrote the manuscript.

## ACKNOWLEDGMENTS

This work was supported by a grant from the National Science Foundation (MCB1411482 to CFL), the National Institutes of Health (U01CA215845 to CFL), The Incyte-Vanderbilt Research Alliance (M. Savona PI, CFL Co-I), and the Oak Ridge Leadership Computing Facility at the Oak Ridge National Laboratory, through the Office of Science of the U.S. Department of Energy under Contract No. DE-AC05-00OR22725.

